# A State Representation for Reinforcement Learning and Decision-Making in the Orbitofrontal Cortex

**DOI:** 10.1101/210591

**Authors:** Nicolas W. Schuck, Robert Wilson, Yael Niv

## Abstract

Despite decades of research, the exact ways in which the orbitofrontal cortex (OFC) influences cognitive function have remained mysterious. Anatomically, the OFC is characterized by remarkably broad connectivity to sensory, limbic and subcortical areas, and functional studies have implicated the OFC in a plethora of functions ranging from facial processing to value-guided choice. Notwithstanding such diversity of findings, much research suggests that one important function of the OFC is to support decision making and reinforcement learning. Here, we describe a novel theory that posits that OFC’s specific role in decision-making is to provide an up-to-date representation of task-related information, called a state representation. This representation reflects a mapping between distinct task states and sensory as well as unobservable information. We summarize evidence supporting the existence of such state representations in rodent and human OFC and argue that forming these state representations provides a crucial scaffold that allows animals to efficiently perform decision making and reinforcement learning in high-dimensional and partially observable environments. Finally, we argue that our theory offers an integrating framework for linking the diversity of functions ascribed to OFC and is in line with its wide ranging connectivity.

## 1. Introduction

The orbitofrontal cortex (OFC) is an intensely studied brain area. Pubmed currently lists over 1,000 publications with the word ‘‘Orbitofrontal” in the title, reflecting more than six decades of research that has mainly sought to answer one question: what mental functions are subserved by the OFC? Still, an integration of existing knowledge about this brain area has proven difficult (Cavada and Schultz, 2000; Stalnaker et al., 2015). Lesion studies have pointed to a plethora of often subtle and complex impairments, but what mental operation is common to all these impairments has remained unclear. Neural recordings have shown that a large variety of different aspects of the current environment are encoded in orbitofrontal activity, but have not yet explained why these variables are co-represented in OFC or how they are integrated. Finally, the study of OFC’s anatomy has uncovered a complex internal organization of sub-regions that has made the identification of homologies in other mammals difficult, and raised questions about the functional division of labor associated with these subregions.

Despite this diversity of findings and the lack of consensus on their interpretation, research has converged on the idea that many of OFC’s functions must lie within the domains of decision making and reinforcement learning. In this chapter, we will provide a selective overview of the studies supporting this broader idea, and describe a novel theoretical framework for understanding the role of the OFC in reinforcement learning and decision making. This framework, the State-Space Theory of OFC, proposes that the OFC represents, at any given time, the specific information needed in order maximize reward on the current task.

We will begin the chapter with a brief overview of OFC anatomy, focusing mostly on its most salient aspects in primates. We will then discuss the State-Space Theory of OFC in more detail and evaluate how well it can accommodate current knowledge. We will follow with a discussion of other major theoretical accounts of the OFC and how they might be integrated into the State-Space Theory. Finally, we will discuss which findings about OFC lie outside the scope of our framework and highlight some areas for future research. Given the large number of investigations, our overview remains necessarily incomplete, and we refer the reader to several recent excellent reviews on this topic for more information (Kringelbach, 2005; Murray et al., 2007; Rudebeck and Murray, 2011; Rushworth et al., 2011; Schoenbaum et al., 2011; Stalnaker et al., 2015).

## 2. What is the Orbitofrontal Cortex?

The primate OFC is a large cortical area located at the most ventral surface of the prefrontal cortex, directly above the orbit of the eyes (hence the name), and including parts of the medial wall between the hemispheres (see Figure 1A). It is defined as the part of prefrontal cortex that receives input from the medial magnocellular nucleus of the mediodorsal thalamus (Fuster, 1997), and consists of Brodman areas 10, 11 and 47. Brodman’s initial classification, however, was unfinished and showed inconsistencies between humans and primates, possibly reflecting the heterogeneity of sulcal folding patterns in OFC (Kringelbach, 2005). Later cytoarchitectonic work has refined this classification and today’s widely accepted parcellations are based on five subdivisions known as Walker’s areas 10, 11, 47/12, 13 and 14 (e.g., Öngür et al., 2003; Glasser et al., 2016, see Fig. 1B, C). One unusual aspect of the primate OFC is its mixed cytoarchitecture that is partly five-layered (agranular) and partly six-layered (granular), unlike the homogenous six-layered structure found elsewhere in the prefrontal cortex. This suggests that OFC is phlyogenetically older than other parts of frontal cortex and complicates comparisons between primates and nonprimates whose OFC is entirely agranular (Preuss, 1995). Based on these differences, Wise and colleagues have proposed that non-primate mammals have no homologue of what is the granular OFC in primates (Wise, 2008). Many controversies about this topic are still ongoing, for instance about whether the medial OFC network does or does not have a homologue in rodents (Schoenbaum et al., 2006; Heilbronner et al., 2016; Rudebeck and Murray, 2011).

**Figure 1:**
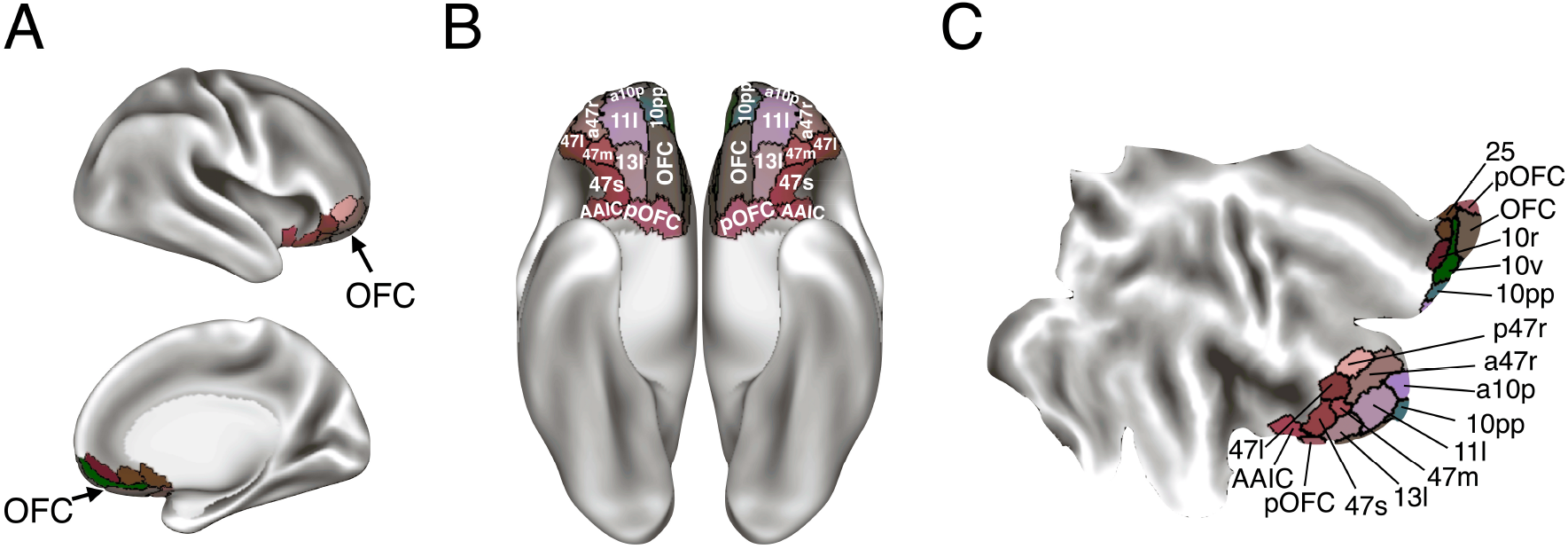
Anatomy of the human orbitofrontal cortex. *(A)*: The location of the OFC on a inflated brain is highlighted by the brown shaded areas in a lateral (upper) and medial (lower) view. *(B)*: Subdivisions of the OFC, shown on the ventral surface, according to most recent parcellation proposed by Glasser et al. (2016). *(C)*: OFC subareas shown on a flat map of the left hemisphere, with the same color coding as in (A) and (B). AAIC: Anterior agranuar insular complex. Figure made using data from Glasser et al. (2016) and the Connectome Workbench.

Another noteworthy feature of OFC’s anatomy is its connectivity. OFC has remarkably close connections to all sensory areas (often only bi- or trisynaptic) in addition to widespread connections to other parts of frontal cortex, striatum, amygdala and hippocampus, amongst others. Connectivity patterns to these areas highlight a distinction between a medial and a lateral sub-networks in the OFC (Cavada et al., 2000; Kahnt et al., 2012), a difference that is often assumed to have functional implications (Rudebeck and Murray, 2011; Noonan et al., 2010b; Walton et al., 2011; Elliott et al., 2000). Specifically, the lateral network has been shown to have many connections to lateral orbital areas as well as the amygdala and receives connections from sensory areas related to olfactory, gustatory, visual, somatic/sensory and visceral processing. The medial network, in contrast, connects to areas along the medial wall (Brodman areas 25, 24 & 32) and receives input from the amygdala, the mediodorsal thalamus, various regions in the medial temporal lobe (hippocampus, parahippocampus, rhinal cortex), ventral striatum, hypothalamus and periaqueductal grey (Cavada et al., 2000). For the remainder of this chapter, we will consider both networks as OFC, following definitions of the orbital medial prefrontal cortex (Öngür et al., 2003). Note that the area commonly referred to as ventromedial PFC is therefore partly included in our definition of OFC.

In summary, the orbitofrontal cortex represents a remarkably densely connected brain area with links to all sensory domains, learning and memory structures like striatum, amygdala and hippocampus as well as several frontal subregions. The OFC is also a highly heterogenous brain area, containing two broadly distinct sub-networks (medial vs. lateral), cytoarchitectonic diverse subregions (granular vs. agranular cortex) and large inter-individual differences in sulcal folding patterns.

## 3. The State-Space Theory of OFC

At the heart of studying decision making is the quest to understand how the brain answers the following question: Given the state of the world, which actions promise to yield the best outcomes? Much research has focused on the only aspect of this question, namely how actions are selected and how expected outcomes are learned and represented. In contrast, the other major aspect of the question, how decisions depend on the current environment, and what the animal considers “the state of the world,” have received much less attention. Because natural environments of animals are often rich in sensory information and complex temporal dependencies, how the state of the environment is represented in the brain is crucial for successful decision making. Below we will define more precisely what we mean by “state of the world,” and provide more detail on how the representation of the state of the world is shaped by the requirements of the decision making process. Then we will propose a specific role for the OFC in representing this information during decision making.

The computational theory of reinforcement learning (RL, Sutton and Barto, 1998) relies on a representation of all the information that is relevant for the current decision, referred to as *the state.* The state is not just a one-to-one reflection of the physical state of the environment, but rather a reflection of what information about the world the decision-making agent represents at the moment the decision is made. How precisely should an agent represent its environment to optimally support decision making and learning? As we will see below, this is not an easy question, and answering it requires a good understanding of the problem at hand. Consider, for instance, an RL agent trying to learn to balance a pole hinged to a cart that can be moved either left or right (a classic benchmark task in RL, Michie and Chambers, 1968). From an RL perspective, an optimal policy for this problem can only be computed if the current state contains all information that is sufficient to fully predict the immediate future state of the cart, if a certain action is taken. This characteristic is known as the *Markovian* property and effectively means that the conditional probabilities of future states depend only on the current state and action, but not the past states. In the case of the pole problem, this means that representing the cart’s position and the angle of the pole as the state are not enough, because these variables alone are insufficient to predict which way the pole is moving and to infer how to move the cart. Instead, the cart’s velocity and the rate at which the angle between the pole and the cart is changing are needed. Representing these variables as part of the current state will allow one to learn a much better behavioral control policy than if they were omitted from the state representation.

This requirement for the state representation raises another problem: some variables do not have a one-to-one correspondence to the information the agent gets from its sensors. For example, the velocity related variables must be inferred by comparing past and current sensory inputs, and thus require memory. If states need to reflect information beyond what is accessible through current sensory input and there is uncertainty regarding their true underlying value, the states are called *partially observable.*

Finally, not all aspects of the current sensory input are relevant. Lighting conditions, for instance, do not need to be included into the state as they are irrelevant for the policy even if they change the sensory signals. Including unnecessary aspects in the state representation will lead to slower learning due to the need to separately learn a policy for states that seem different but are effectively equivalent, a phenomenon known as *the curse of dimensionality.* A good state representation is therefore one that solves two problems: it deals with partial observability and non-Markovian environments by supplementing sensory information with the necessary unobservables, and it filters the sensory input to only include relevant aspects in order to avoid the curse of dimensionality.

While pole balancing itself is a rare activity for humans, the curse of dimensionality and the problem of partial observability of states are ubiquitous. A brain area well suited to solve this problem would need to be able to access sensory cortices as well brain areas relevant for episodic memory and selective attention processes (Niv and Langdon, 2016). On purely anatomical grounds, the OFC is a good candidate for this representation: it is unique among areas in the prefrontal cortex in its close connectivity to all five sensory modalities, and it has bidirectional connections to brain areas relevant for memory and decision making such as the hippocampus and the striatum. In addition to these general considerations, a review of decades of studies on the function of the OFC has recently led us to propose that the role of the OFC in decision making is to represent partially observable states of the environment when they are needed to perform the task at hand. Specifically, in Wilson et al. (2014), we investigated how changes in the way states are represented would affect behavior in tasks that are known to be impacted by OFC lesions. The central idea was that in many cases the state space of a task must include partially observable information, but OFC lesioned animals might be incapable of integrating the necessary observable and unobservable information. In order to test this idea theoretically, we used an RL modeling framework and manipulated how states were represented. To simulate OFC lesioned animals, all states of the task that are associated with identical sensory input were therefore modeled as the same state, whereas healthy animals were modeled as having the ability to disambiguate states that involve identical sensory input based, for instance, on past events. Strikingly, this manipulation caused subtle but pervasive impairments of the model’s ability to perform exactly those tasks that are known to be impacted by OFC lesions.

One example is the delayed alternation task, which is known to be impaired by OFC lesions in both animals as well as humans (e.g., Mishkin et al., 1969; Freedman et al., 1998). In this task, two simple actions (say, pressing a left or right lever) can lead to reward. Specifically, the delivery of reward is coupled to the previous choice such that on each trial, only the action that was *not* chosen on the previous trial is rewarded. To solve this task, the states corresponding to the two options need to be supplemented by the previous choice: although all trials look similar in terms of the externally available stimuli, when the previous choice was A, the best action is B, and vice versa (see Fig. 2A/B). If OFC lesions impair the ability to distinguish between two identically-looking states based on unobservable context, one would expect that OFC-lesioned animals would represent the task as having only one state (Fig. 2B). As a consequence, the animal would be severely impaired in its ability to correctly perform the task, which indeed has been shown (specifically, performance went down to chance due to the lesion, but only if trials were separated by a delay that rendered the previous choice unobservable; Mishkin et al., 1969). In Wilson et al. (2014), we showed that a variety of behavioral consequences of OFC lesions can indeed be explained by an impairment in the state space underlying performance on the task. We also showed that changes in dopaminergic firing following OFC lesions can be explained as a consequence of impaired state differentiation (Takahashi et al., 2011).

**Figure 2:**
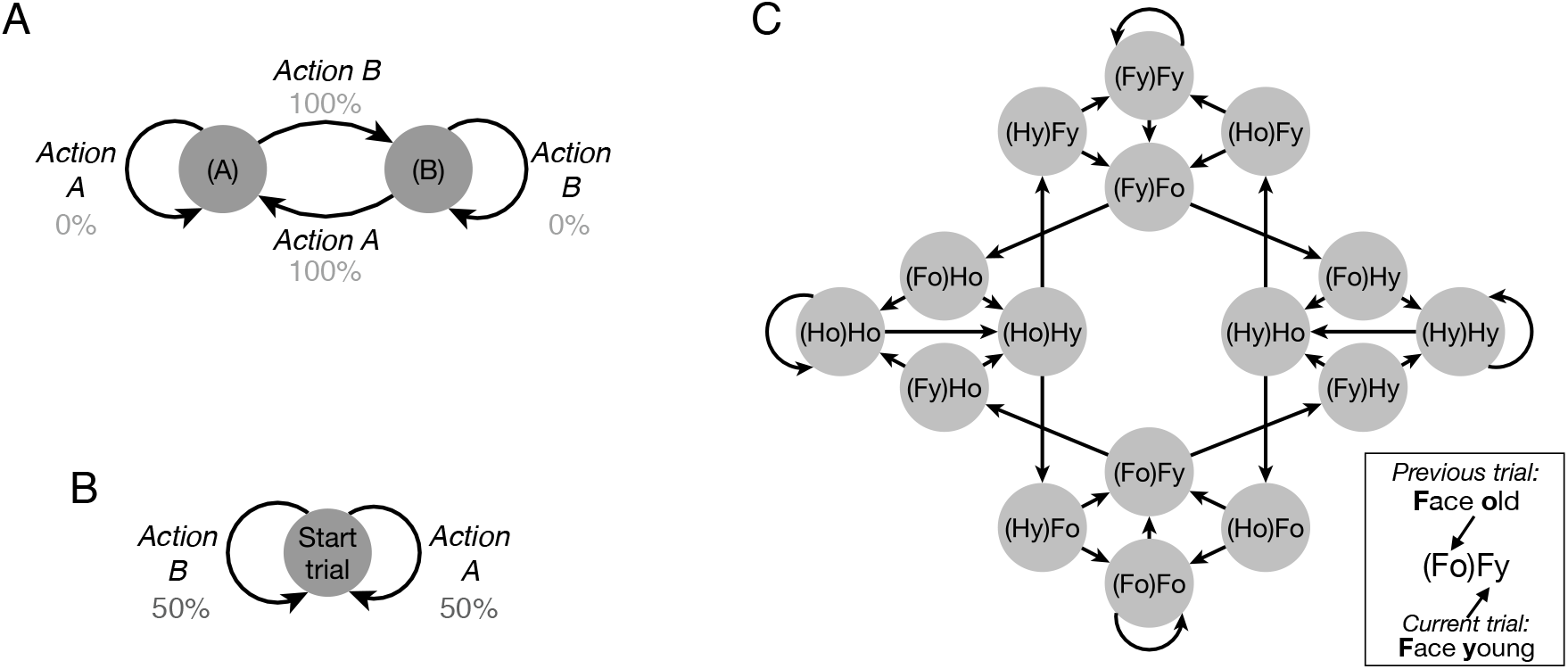
State spaces of the delayed alternation task and the Schuck et al. task. *(A)*: In a delayed alternation task, performance is rewarded if the previously unchosen action is selected. A suitable state representation therefore must distinguish between trials in which action A was previously performed versus trials in which action B was previously taken. We denote the two possible actions as ‘A’ and ‘B’ and label the states accordingly (A) and (B) to denote the action on the previous trial, see Fig. 2A. In the diagram, state transitions depend on the action taken. The probability of reward on each trial, with depends on the transition/action chosen, is denoted in gray for each transition. With such a state representation, different values can be assigned to an action depending on whether is was preceded by the same or a different action, thus allowing the agent to learn the optimal policy that leads to 100% reward (alternating choices). *(B)*: If, on the other hand, an agent is unable to differentiate between states based on the unobservable choice history, then the environment is perceived as having only one state, in which each action yields reward on only 50% of trials. According to the theory, complete OFC lesions would result in this reduced state representation, and consequently performance would be at chance accuracy, as is indeed seen empirically (Mishkin et al., 1969). *(C)*: The state space used in the Schuck et al. task described in the text. ‘Hy’ indicates a trial in which the relevant category was House, and the correct response was young. For simplicity, only transitions for correct actions are shown (the wrong action leads to repetition of the trial). State-relevant information from the previous trial is denoted in brackets, such that ‘(Fo)Fy’ indicates a young Face trial (the ‘Fy’ part of the state) that was preceded by an old Face trial (the ‘(Fo)’ part of the state, see legend).

In a follow up study in humans, we used a task specifically designed to test our hypothesis, to investigate orbitofrontal representations during decision making (Schuck et al., 2016). On each trial of the task, participants had to judge whether either a face or a house (presented overlaid as a compound stimulus) was old or young. Crucially, to determine whether they should be judging the house or the face, participants had to continuously monitor both the current and previous trial: whenever the age response on the previous trial was different from that on the current trial, the category to be judged on the following trial was switched. Otherwise, the next trial’s category was the same as the current trial. Given these rules, the task required a complex state space with 16 different partially observable states (Fig. 2C). Using multivariate pattern analysis (MVPA) techniques, we investigated what aspects of the state information were encoded within OFC. This analysis showed that on each trial, the OFC contained information about all partially observable aspects of the state: the previous age, the previous category and the current category. A whole-brain analysis suggested that medial OFC was the only region in which all necessary unobservable information could be decoded. Still, information about events two trials in the past, which was not relevant to correct performance, could not be decoded in OFC (in contrast to other brain areas where we could decode only some of the relevant unobservable state components, but also some irrelevant information such as the category from two trials back). Finally, in an error-locked analysis of single-trial information, we found that errors during the task were preceded by a deterioration of the state representation in OFC. These results provide strong support for the above outlined hypothesis of the representational role of OFC in decision making.

Several other studies have come to similar conclusions. Recording from neurons in lateral orbitofrontal cortex in rodents, Nogueira et al. (2017) reported that task-relevant but unobservable information from the previous trial was integrated with the current sensory input. A study from our lab used a task in which participants had to infer the current state based on a series of past observations, and found that activity patterns in OFC reflect the posterior probability distribution over unobserved states, given the observed sequence of events (Chan et al., 2016). Bradfield et al. (2015) reported that bilateral excitotoxic lesions and designer-drug induced inactivations of the rat medial OFC led to an inability to retrieve or anticipate unobservable outcomes across a range of tasks. Finally, Stalnaker et al. (2016) studied rats in a task in which outcome magnitudes and identities were occasionally reversed. They found that the unobservable state of the task (that is, the “block identity”) could be decoded from activity of cholinergic interneurons in the dorsomedial striatum, and importantly, that this information vanished when the OFC was lesioned. Taken together, these studies support the idea that within OFC, task-relevant information is combined into a state representation that facilitates efficient decision making in the face of partial observability and non-Markovian environments.

## 4. A role for the OFC in state inference and belief states?

If the OFC is involved in representing partially observable information in the service of decision making, one important question is whether OFC is also involved in *inferring* the state from observations. As we highlighted above, a useful state representation is not simply a reflection of the current sensory input, but rather can be viewed as the (often hidden) collection of attributes that causally determine future rewards and state transitions. For example, in the young/old task described above, the current stimulus is not sufficient for determining which action will be rewarded. Similarly, when deciding whether the cab you requested still on its way, or have they forgotten your request and you should call the company again, the current observation of “no cab here” is not sufficient and you must make use of information such as how long you have already been waiting, what is the time, and what time did you request the cab for. This implies that the current state must often be *inferred* from more than current observations, and in many cases there is considerable uncertainty about the current state. Reinforcement learning theory has shown that an optimal way to learn under such uncertainty is to use Bayesian inference to estimate the probability distribution over possible unobservable states given the observations, and use this quantity (referred to as the *belief state*) as the current state of the task (Kaelbling et al., 1996; Rodriguez et al., 1999; Daw et al., 2006; Dayan and Daw, 2008; Rao, 2010; Samejima and Doya, 2007).

Although it is still unclear whether the brain indeed performs a similar inference process and if the OFC is representing a belief state distribution rather than a single (for instance, most likely) state, recent evidence has pointed in that direction. For example, in a recent study, we studied the process of inferring the true state in a task in which observations were only probabilistically related to states (Chan et al., 2016). Using a representational similarity approach, we found that the similarity of neural patterns in medial OFC was related to the similarity of probabilistic state distributions predicted by a Bayesian inference model. In line with this idea, other studies have indicated that OFC also represents the confidence with which animals make a choice (Kepecs et al., 2008; Lak et al., 2014), a quantity that also affects dopaminergic midbrain activity and may reflect belief states (Lak et al., 2017). Other studies investigating dopaminergic prediction error signals have shown that reward predictions are based on a state inference process rather than purely sensory states (Starkweather et al., 2017; Langdon et al.). In particular, the passage of time provides a ubiquitous cue for inferring transitions between states that may be externally similar (as in the cab example above), and recent work suggests that prediction error signals rely on input from the ventral striatum to reflect such time-based state inference (Takahashi et al., 2016). Our previous work has shown that dopaminergic prediction errors indeed depend on state representations in the OFC (Wilson et al., 2014; Takahashi et al., 2011). Together with this recent work that indirectly suggests the existence of belief states elsewhere in the brain, these findings support the idea that a state inference process results in a representation of an entire belief-state distribution within the OFC. More evidence is needed, however, and previous work has suggested that belief states may be encoded either in lateral PFC (Samejima and Doya, 2007) or sensory cortices (Daw et al., 2006; van Bergen et al., 2015).

In addition, many questions about the inference process itself remain unanswered. Of particular importance is the question about the role of state transitions in the state inference process. The above mentioned theoretical approaches to belief states indicate that inferring the current belief state relies on two quantities: current and past observations on the one hand, and the previous belief state on the other hand (see Figure 3). Previous belief states influence the current belief state through the state transition function: similar to how knowledge about someone’s previous location will constrain where that person could possibly be one time step later, knowledge about possible state transitions and the previous state will influence the estimate of the current state. Several of the tasks in which OFC lesions lead to behavioral impairments, such as devaluation (Gallagher et al., 1999; Pickens et al., 2003) and Pavlovian-instrumental transfer (Bradfield et al., 2015) tasks, not only require state representations, but also inference about expected outcomes based on knowledge of state transitions. Furthermore, neuroimaging studies have found that the OFC may be involved in updating knowledge about transitions between cues and outcome identities (Boorman et al., 2016), which could reflect a role in representing state transitions more generally. Other studies have pointed to a prominent role of the hippocampus in storing state transitions (Schapiro et al., 2012).

**Figure 3:**
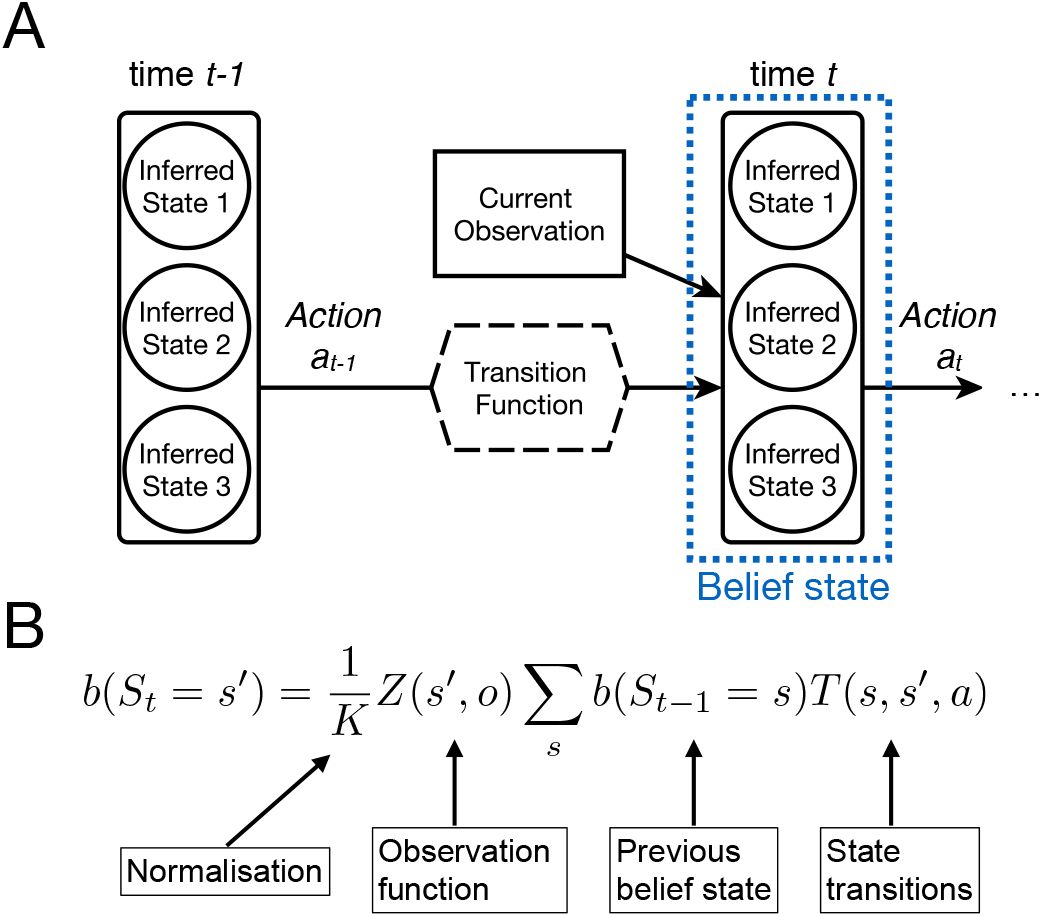
Belief state representations and state inference. *(A)*: The diagram illustrates how uncertainty at different stages is incorporated into a belief state representation (blue box). On each trial, because of partial observability, we assume that the sensory input (‘Current Observation’) cannot be deterministically mapped onto states, but rather leads to a probabilistic estimate of how likely different states are (captured by the *observation function Z*(*s′,o*)). In addition, the distribution over states on the previous trial is combined with the (presumably known) state transition function to further constrain the likely current state. In analogy to a spatial setting, this process corresponds to estimating your current location based on what is currently observed, in combination with your previous belief regarding where you have just been, which determines the locations that were adjacent to you and reachable in one step. Together, the current sensory input and your previous beliefs regarding your likely location constrain your current location estimate. *(B)*: The key belief-state updating equation relates the above described quantities to yield the probability of being in a specific state *s*’ at a specific time *t*, denoted as *b*(*S_t_* = *s*′).

Finally, an important aspect of state inference is that the environmental features that are predictive of the outcome might change over time, or previously unknown relations between sensory input and outcomes might only be discovered after a period of learning. In our previous work, we have begun to investigate both of these cases (Schuck et al., 2015; Niv et al., 2015). This work has so far pointed not towards the OFC, but rather suggested a role for a frontoparietal attention network in controlling the adaptation of state representations, and of medial prefrontal cortex in covertly preparing the updated state representations.

In summary, current research bears only indirectly on the possible role of OFC in the representation of belief states, state transitions and state updating processes and it seems unlikely that all these functions would be performed by a single neural circuit. Future research therefore needs to address these questions more directly, and also investigate how different brain areas such as the hippocampus and medial prefrontal cortex might perform these complex computations in cooperation with the OFC (e.g., Kaplan et al., 2017). Moreover, one important question is where and how the relations between states, as reflected in the possible transitions between them, are represented in the brain. Building on the existing work about hippocampal representations of transitions between spatial as well as observable states (Schapiro et al., 2012; Stachenfeld et al., 2017), future work needs to investigate the neural representation of transitions between partially observable states. Such research needs to carefully take into account the methodological hurdles of estimating representational similarity with fMRI, however (Cai et al., 2016).

## 5. Orbitofrontal value signals and their role in decision making

While the work reviewed in the previous sections provides support for the State-Space Theory of OFC, these findings are only a small part of the large literature on the OFC, and other authors have proposed different accounts of OFC function. Perhaps the most influential alternative is that OFC represents economic value associated with a given choice, in particular value that has to be calculated on-the-fly rather than learned from experience. This theory has its roots in recordings of neural activity in the OFC of monkeys making choices between different food options (Tremblay and Schultz, 1999; Padoa-Schioppa and Assad, 2006). In particular, in one influential study, Padoa-Schioppa and Assad (2006) recorded activity in the OFC (area 13) of monkeys while the monkeys chose between two different kinds of juice. On each trial, different amounts of each juice were offered through the display of visual cues, and the animals could freely decide which option they preferred. Following standard economic theory, the authors calculated the subjective value of each juice based on the monkeys’ choices and showed that a proportion of recorded neurons showed firing activity that varied linearly (some increasing and some decreasing) with the subjective value of the chosen juice, regardless of which option was chosen (they additionally reported neurons that responded to the identity of the chosen option, and the value of each of the two offers). This general finding, a value or reward representation that is independent of sensory or motor aspects of the option, is supported by a number of similar studies in humans, monkeys and rodents (e.g., Thorpe et al., 1983; Schoenbaum and Eichenbaum, 1995; Tremblay and Schultz, 1999; Gottfried et al., 2003; Plassmann et al., 2007; Hare et al., 2008; Howard et al., 2015). Moreover, subsequent studies have suggested that these value representations do exhibit a number of properties in line the value account of OFC function. For example, Padoa-Schioppa and Assad (2008), found that OFC value signals are invariant to the other available options (a property called transitivity; but see Tremblay and Schultz, 1999).

What these findings imply about OFC function is not as clear as it might seem, however. First, due to the long recording durations and many experimental sessions involved in monkey electrophysiology, it is difficult to claim with certainty that the monkeys are computing the values of the alternatives on the fly, and that this is what OFC is necessary for. Moreover, other findings have cast doubt on the claim that OFC’s primary function is to represent values of choices during decision making in general (Schoenbaum et al., 2011). For example, studies of OFC-lesioned animals have shown no impairment in general learning abilities during initial value acquisition (e.g. Butter, 1969; Chudasama and Robbins, 2003; O’Doherty et al., 2003a; Izquierdo et al., 2004; Chudasama et al., 2007; West et al., 2011), and inconsistent results after a reversal of cue-outcome contingencies (Kazama and Bachevalier, 2009; Rudebeck et al., 2013; Stalnaker et al., 2007). Other studies have shown that OFC lesions do not impact monkeys’ ability to make value-dependent choices even when values are constantly changing (Walton et al., 2010). A recent proposal by Walton et al. (2011) has highlighted that the nature of the decision-making impairment depends on whether the lesion affected medial or lateral OFC. Specifically, Walton and colleagues suggested that lateral OFC is necessary for correctly assigning credit for outcomes to previous actions for the purpose of learning, whereas the medial OFC is important for basing decisions on the highest-valued option while ignoring the irrelevant other options. This idea might indeed explain why decision making impairments are seen under some circumstances but not others.

Yet, values (and choices) are not the only task-related quantity that is encoded in OFC. Several reports have shown that OFC encodes many variables that are related to the current task but are independent of value, including for instance outcome identities (McDannald et al., 2011, 2014; Howard et al., 2015; Stalnaker et al., 2014), salience (Ogawa et al., 2013), confidence signals (Lak et al., 2014; Kepecs et al., 2008) and even social category of faces (Watson and Platt, 2012) or spatial context (Farovik et al., 2015). In fact, these value-free signals may represent the vast majority of information encoded in OFC - one study found only 8% of lateral OFC neurons coding value in a linear manner (Lopatina et al., 2015), a number that is not out of line with other reports that often analyze only small subsets of the recorded neurons. This raises the possibility that firing patterns of orbitofrontal neurons reflect value only in the context of, or as part of, the current state. This interpretation is also supported by the above-described study, in which we found state signals in the OFC in the absence of any overt rewards or values (Schuck et al., 2016).

This idea that OFC value representations are embedded in a more general state signal in the OFC is supported by a recent study from Rich and Wallis (2016). In this experiment, OFC neurons were recorded while monkeys deliberated between two choices leading to differently valued outcomes. While activity patterns during deliberation were predictive of the value of the later chosen option, the authors also report that the value encoded by single neurons was dependent on a network-represented state: the same neuron encoded the value of pictures on the right when the rest of the network (not including this neuron) was in the ‘right state” (i.e., signalling that something was shown on the right), and the value of pictures on the left when the network was in the “left state.”

The notion that the unique function of OFC is to integrate information about partially observable task states (and perhaps their resulting values) is also supported by research on flexible, goal directed behavior in the so-called devaluation paradigm. In this task, animals are first trained to perform actions (say, pressing on one of several bars) in order to obtain desired outcomes (e.g., different types of food) such that each action is associated with a particular outcome. Subsequently, in a separate setting, one of the possible outcomes is devalued by satiation or pairing the outcome with food poison. A test then assesses whether the animal can use the new (lower) value of the outcome to guide its behavior. In this, the animal is once again allowed to make outcome-earning actions (although no outcomes are actually delivered in this phase), with the main question being whether it will continue to perform the action that previously led to the now devalued outcome. While healthy animals that have not been over-trained to the point that the actions have become habitual adapt their behavior appropriately, OFC-lesioned animals continue to chose the wrong option, a behavior that has been suggested to reflect their inability to update outcome expectations (Gallagher et al., 1999; Pickens et al., 2003), rather than mere failures to inhibit prepotent reposes (Chudasama et al., 2007; Walton et al., 2010). In another devaluation study, value representations in the OFC as well as the Amygdala were found to be changed following devaluation (Gottfried et al., 2003), suggesting that the values encoded in OFC can be updated offline based on knowledge of state transitions and the newly experienced outcomes. Interestingly, consistent with the State-Space Theory, similar studies looking at the representations of *value-independent* outcome identities in the OFC found that these representations differed based on the current goal, consistent with the state-space representation being task dependent (Critchley and Rolls, 1996; Howard and Kahnt, 2017). Thus, the role that OFC seems to play in value-based decision making seems to be at the junction of representing values and correctly inferring partially observable current or future states.

Other studies using unblocking paradigms support similar conclusions (McDannald et al., 2011; Burke et al., 2008). In one variant of this task, animals learn to discriminate different odors that predict different quantities and flavors of milk. After learning, additional odors are added to the original odor and either the same outcome is presented, the size of the outcome is changed, or the flavor or the outcome is changed. Afterwards, the degree to which an association between the novel odors and the outcomes was learned is assessed. Because value-based learning is driven by the difference between one’s expectations and the outcomes (the so-called prediction error), learning theory suggests that no association should be learned for the novel odor when it led to the same outcome as before (hence the term “blocking” as the association between the old odor and the outcome blocks a new association from forming Kamin, 1969). The novel odors associated with a change in reward size, however, should be followed by a prediction error that would trigger learning (“unblocking”). This learning can be based purely on value signals, as found in areas such as the striatum. A very different and interesting prediction arises when considering trials in which the milk flavor was changed. Because the two milk flavors were matched for overall value, learning about novel cues predicting flavor changes cannot be driven by a value mismatch error. Rather, for changes in the outcome identity to trigger learning about the new odor specific knowledge about the expected identity of the outcome (and its violation) is required, as found in the OFC (Critchley and Rolls, 1996; Howard and Kahnt, 2017; McDannald et al., 2014; Stalnaker et al., 2014). Indeed, lesion studies have shown that OFC is critical for this type of outcome-identity dependent unblocking but not for unblocking due to changes in the value of the outcome in this task (McDannald et al., 2011).

In summary, while these findings suggest that orbitofrontal neurons encode information about the values of different options, OFC’s function is not readily captured by the proposal that the sole or primary role of this area is value comparisons. From the perspective of the State-Space Theory, these findings rather point to a more holistic integration of decision-relevant information in the OFC, ranging from the potentially partially observable context that is necessary for solving the task, to the expected sensory aspects of the outcomes and the values associated with them. Interestingly, the above described lesion and inactivation studies all suggest that this representation is only necessary when changes in the contingencies between states require to reassess the value of different choices.

## 6. Beyond learning and decision making

Although the literature on the function of the OFC has mainly addressed its role in decision making, some investigations have focused on other potentially important aspects of OFC function. In particular, observations from human patients with OFC damage have often led to reports about post-lesion changes in their ‘personality’ (e.g., Galleguillos et al., 2011; Cicerone and Tanenbaum, 1997). These clinical impressions are corroborated by studies that have established links between between OFC and aggressive behavior (e.g. Raleigh et al., 1979; Butter et al., 1970; Beyer et al., 2015), processing of social (e.g. O’Doherty et al., 2003b; Ishai, 2007; Perry et al., 2016; Azzi et al., 2012) or emotional information (e.g. Schutter and van Honk, 2006; Kumfor et al., 2013; Bechara, 2004; Izquierdo et al., 2005), and risky or impulsive behavior (Bechara et al., 2000). In addition, some studies have indicated that OFC may be important for long-term memory (Meunier et al., 1997; Frey and Petrides, 2002; Petrides, 2007) and working memory (Barbey et al., 2011).

Given the complex anatomy of the OFC, such diversity of findings is not surprising. Previous research has shown that lesion effects can be the result of damage to passing fibers rather than damage to OFC per se (Rudebeck et al., 2013). Moreover, some effects may result from co-damage to other areas (Noonan et al., 2010a), and incorporating the effects of connected areas is generally an important approach for understanding OFC’s function (Rempel-Clower, 2007). In addition, it is certainly possible that some subregions of OFC have functions outside of the domain of decision making.

While our framework does not aim to account for the entirety of the function of this large brain area, it is noteworthy that some of the effects mentioned above are not orthogonal to our proposal. Studies investigating the processing of emotional and facial information have used stimuli that could also be interpreted as either having positive (smiling, attractiveness) or negative value (angry facial expression) (Chien et al., 2016; Winston et al., 2007). In general, inferring others’ emotions and intent are the epitome of inference of a partially observable state. Indeed, these tasks require participants to process “subtle social and emotional cues required for the appropriate interpretation of events” (Cicerone and Tanenbaum, 1997, abstract), which could be affected by lesioned patients’ inability to integrate partially observable information and current sensory input into a suitable state representation. Moreover, the reported association with working memory seemed to be specific to n-back tasks (Barbey et al., 2011) that also require decision making and are similar to the task we used to test our theory in humans (Schuck et al., 2016).

Apart from these issues of integrating all available evidence, one important avenue for future research is to specify predictions about the OFC representations that drive decision making in different tasks and how these OFC representations are affected by lesions and disease. Existing computational models therefore need to be specified in order to predict which hidden and observable aspects of the environment need to be incorporated into the current state in order to solve the task sufficiently (see Collins, Chapter 5 in this book). Ideally, models and theory would also yield predictions about how these state representations develop during task learning, depending on the specific history of choices and experiences of each subject (Niv et al., 2015; Gershman et al., 2015).

## 7. Summary

In this review, we have focused on work that links the OFC to many aspects of decision making. Several prominent findings have reported that individual neurons in OFC linearly increase their firing with the value of the chosen option. Other studies, however, have highlighted that many different aspects of the ongoing task are encoded in OFC’s neural activity, that value neurons dynamically change which option they are encoding depending on the network state, and that value neurons make up only a small proportion of the OFC population in certain circumstances. Moreover, studies of OFC lesions are not consistent with the claim that the OFC is the brain’s sole site of performing basic value-based decision making. Rather, behavioral impairments seem to occur only under certain circumstances, e.g., when a change of previously learned associations is prompted by drastic changes in outcome value, when task rules require rapid switching between values, or when learning must focus on the sensory properties of the outcome rather than its value. In all these cases, performance depends critically on the ability to *infer* a partially observable state of the environment, and maladaptive behavior becomes visible when the state changes in the absence of any sensory cues, and previously learned values no longer apply.

We therefore suggest that the role of the OFC in decision making is to represent the current state of the environment, in particular if that state is partially observable. We have presented several lines of evidence, including lesion studies, electrophysiological recordings, computational modeling and fMRI, that support our framework. We also outlined avenues for future research that should seek to directly investigate to what extent previously reported associations between OFC and cognitive functions outside of the domain of decision making could also result from changes in partially observable state representations (e.g., working memory or personality changes). In addition, taking into account anatomical diversity within OFC and, in the case of lesion studies, careful scrutiny of the nature of OFC insults (e.g., if passing fibers have been damaged) might clarify the origins of some of the diversity of functions associated with the OFC. Crucially, we argued that computational models that specify the representations underlying successful decision making need to be advanced, and tested against empirical data. Together with the existing evidence, these efforts promise to yield unprecedented insight into the functions of an elusive brain area.

## 8. Acknowledgements

This work was supported by the grant 1R01DA042065 from the National Institution on Drug Abuse at NIH awarded to YN. We thank Matthew Glasser for advice in making the OFC map shown in Fig. 1, and Sam Chien for feedback on this manuscript.

